# Sperm-specific fertility factors SMZ-1/2 promote FB-MO biogenesis and MSP filament assembly in *C. elegans*

**DOI:** 10.64898/2025.12.01.691703

**Authors:** Hsiao-Fang Peng, Chih-Ling Liu, Wann-Neng Jane, Chang-Shi Chen, Chao-Wen Wang, Jui-ching Wu

## Abstract

In nematode sperm, motility is powered by polymerization of major sperm protein (MSP) into filaments assembled from a specialized organelle (FB–MO), but how filament formation begins is unclear. We identify two sperm-specific PDZ proteins, SMZ-1 and SMZ-2 (SMZ-1/2), that initiate MSP loading into pre-formed membranous organelles in Caenorhabditis elegans. SMZ-1/2 were expressed specifically in spermatogenic germ cells, colocalized with developing FB-MOs from diplotene through spermatid formation, and were redundant for fertility. Loss of SMZ-1/2 abolished MSP filaments, left MOs MSP-negative and morphologically immature, and caused primary spermatocyte arrest. Electron microscopy showed fewer, smaller FB-MOs lacking crystalline FBs, consistent with a defect in assembly initiation rather than late polymerization. SMZ-1/2 cytoplasmic structures were absent in spe-6 casein kinase mutants, placing SMZ-1/2 within a SPE-6-dependent initiation module. These findings identify SMZ-1/2 as PDZ scaffold proteins that initiate MSP assembly and couple kinase signaling to FB-MO biogenesis during nematode spermatogenesis.

## Introduction

Sperm motility is crucial for reproductive success across animals [1–3]. Following male meiotic divisions, haploid spermatids undergo spermiogenesis, during which they remodel to acquire a cytoskeleton-based motility apparatus. Unlike the flagellated sperm of many species, nematode sperm, including *C. elegans*, exhibit amoeboid motility driven by the actin-like polymerization of major sperm protein (MSP) into filaments that generate force [4, 5]. MSP is essential not only for sperm motility but also for broader aspects of male fertility, functioning as a signaling molecule that induces ovulation [6, 7]. MSP is encoded by multiple gene family and comprises up to 15% of the total sperm protein content [8–10]; thus, proper storage and timely activation of MSP filament assembly are critical for reproductive success. Supporting this, male worms with premature MSP filament activation display impaired male fertility [11].

Early studies showed that MSP is synthesized during spermatogenesis and packaged into a specialized membranous organelle derived from the endoplasmic reticulum and Golgi apparatus [12]. Within this compartment, MSP polymerizes into paracrystalline fibrous bodies (FBs) that become embedded in the membranous organelle (MO) to form the FB-MO complex, a storage site for MSP prior to sperm activation [3, 12, 13]. Genetic analyses have identified key regulators of FB–MO formation. The lysosomal trafficking factor SPE-39 and the presenilin family protein SPE-4 promote MO formation [14–16]. As MOs form, MSP filaments begin to assemble adjacent to the developing MO and gradually extend to form FB. During FB elongation, the intrinsically disordered protein SPE-18 localizes to the ends of the FB, where it recruits MSP into MOs and promotes FB growth [17]. Because FB-MOs are later mobilized during sperm activation to release zinc and drive MSP-based motility, their proper assembly is critical for successful fertilization [18–22].

It has been shown that the initial assembly of MSP filament requires the casein kinase SPE-6 [23]. Nonetheless, the molecular mechanisms that initiate MSP filament assembly and coordinate FB–MO biogenesis remain poorly understood. Proper signal transduction pathways are likely required to integrate MSP polymerization with FB-MO formation, and scaffold proteins may play key roles in this process. PDZ domains are well-characterized protein–protein interaction modules that recognize C-terminal motifs of target proteins [24, 25]. By organizing signaling complexes, PDZ proteins often function as scaffolds that couple membrane-proximal cues to cytoplasmic factors. In spermatogenesis, PDZ domain proteins have been implicated in processes such as the acrosome reaction in mammals [26–30], suggesting that analogous PDZ-mediated mechanisms may also regulate FB-MO assembly in *C. elegans*.

In search for additional male fertility factors, two sperm-enriched PDZ domain-containing proteins, SMZ-1 and SMZ-2 (Sperm Meiosis PDZ-1/2, collectively SMZ-1/2) were identified through comparative proteomic analysis of *C. elegans* sperm and oocyte chromatin [31]. Although highly expressed in spermatogenic germline, their functional roles in spermatogenesis remained unknown. Here, we demonstrate that SMZ-1 and SMZ-2 are essential for initiating MSP filament assembly. Loss of SMZ-1/2 causes primary spermatocyte arrest and underdeveloped FB-MOs absent of MSP filaments, indicating that these proteins are critical for MSP filament assembly. Given their spatiotemporal association with MSP and FB-MO development, our findings suggest that SMZ-1/2 act as initiators of MSP filament assembly, thereby ensuring proper sperm development in *C. elegans*.

## Material and Method

### Strains and culture

*Caenorhabditis elegans* strains were cultured on NGM plates seeded with *E. coli* OP50 as described [32]. Except the temperature-sensitive strains BA17 *fem-1 (hc17)* and JK816 *fem-3 (q20)* were maintained at 15℃. The *smz-1(tpe9)* allele was generated through CRISPR/Cas9 in this study. Other strains used in this study are listed in Supplementary Table 1.

### CRISPR/Cas9

*smz-1(tpe9[3xFLAG::smz-1])* was generated through ribonucleoprotein (RNP)-based editing co-CRISPR injection into N2 as described by Dokshin et al. [33]. The crRNA was designed and ordered from IDT website and the repair templates were generated through hybridized PCR products from pJJR82 (Addgene) [34]. The sequence of the crRNA and the primers for repair template PCR are provided in Table S1. The insertion was targeted after the start codon of *smz-1* and contained a SpeI enzyme site to facilitate genotyping. The F1 rollers were let egg laid and followed by PCR and SpeI enzyme digestion selection.

### Fertility analysis

#### Hermaphrodite progeny production assay

The self-progeny from N2, *smz-1(tm3228)*, *smz-2(wf336)*, and *smz-1(tm3228); smz-2(wf336)* hermaphrodites were determined by placing single worms on separated culture plates. Worms were transferred into fresh plates twice daily until the worms stopped laying eggs. The total laid eggs were counted immediately after the worms were shifted and the hatched larvae were calculated after 72 hours.

#### Hermaphrodite fertility rescue assay

To identify whether the origin of infertility, L4 virgin hermaphrodites from N2, *smz-1(tm3228)*, *smz-2(wf336)*, and *smz-1(tm3228); smz-2(wf336)* were separately mated with *fog-2* males for 24 hours at 20℃. Then the mated hermaphrodites were singled to culture plates and the total hatch rates were counted for two days.

#### Live imaging

The male worms expressing mCherry-histone were dissected in 3 μl complete sperm medium [50 mM Hepes, 45 mM NaCl, 25 mM KCl, 1 mM MgSO_4_, 5 mM CaCl_2_, 1 mg/ml BSA, PH7.0] to release the gonad. The coverslip covered to the droplet to help the spermatocyte releasing and sealed with Vaseline to prevent evaporation [35, 36]. The images were acquired under 100x oil immersion objective with Olympus IX83 microscope system.

#### Immunohistochemistry

For pH3 and α-tubulin staining, male worms were dissected into 5 μl sperm salts [10 mM Pipes, pH 7, 5 mM KCl, 0.2 mM MgSO4, 9 mM NaCl, and 0.4 mM CaCl2] and fixed with equal volume of 4% paraformaldehyde in sperm salt on a SuperFrost Plus Microscope slide. The slides were incubated at room temperature for 5 minutes, frozen in liquid nitrogen for more than 15 minutes, and the cover slip was moved away quickly to tear the tissue apart [35]. Then the slide was put into ethanol for five minutes, washed with PBST twice for 5 minutes, and PBS for 10 minutes once. Primary antibody was incubated in moist chamber at 4℃ overnight, as secondary antibody was incubated at room temperature for two hours followed by PBST/PBS wash. Sample was mounted with Vectashield containing 2 μg/ml DAPI.

For FB-MO and SMZ staining, the male gonads were released in 5 μl sperm salts. After the gonad dissection, additional 5 μl sperm salts were added to the sample. Then the samples were incubated and freeze/crack as described above. The slides were fixed in cold methanol and cold acetone ten minutes for each at 4℃ and followed by ddH2O rinse and ten minutes PBS wash [31]. Slides were incubated with antibodies and mounted as above.

The following antibodies were used in this study: mouse anti-α-tubulin (DM1α, Sigma, T9026) at 1:200, mouse anti-phospho-histone3 (Millipore, T9026) at 1:200, rabbit anti-SMZ (Gene Script, this study) at 1:50, mouse anti-MSP (4A5 from Hybridoma Bank) at 1:5 dilutions, AlexaFluor 488-conjugated wheat-germ agglutinin at 5 μg/ml (Invitrogen), anti-mouse AlexaFluor 488 (Jackson ImmunoResearch, 115-545-003) at 1:200, anti-rabbit AlexaFluor 488 (Jackson ImmunoResearch, 111-545-003) at 1:200, and anti-rabbit AlexaFluor Cy3 (Jackson ImmunoResearch, 111-165-003) at 1:200 dilutions. The staining results were visualized with Zeiss LSM880 confocal microscopy Airyscan under 40X and 100X oil objective and Zeiss LSM980 confocal microscopy under 63X oil objective. The images were analyzed with ImageJ Fiji and were processed with both Adobe Photoshop and Adobe Illustrator 2025 software. The 3D reconstruction images were applied and reconstructed with Imaris (Oxford).

#### Western blot

Ten gravid *fem-1(hc17)* and *fem-3(q20)* hermaphrodites were allowed to lay eggs for 8 hours on fresh NMG plates, then removed. Progeny were grown to adult stage for three days at 25℃ to induce phenotypes. To collect male worms, five L4 *him-5(e1490)*, *smz-1(tm3228)/ mIs11; smz-2(wf336)* hermaphrodites were singled to NGM plates and permitted to lay eggs for four days at 20℃. Hand-picked 100 *fem-1(hc17)*, *fem-3(q20)* hermaphrodites and *him-5(e1490)*, *smz-1(tm3228); smz-2(wf336)* males were collected in 10 μl M9 buffer. Then quickly freeze and thawed for five times in liquid nitrogen to crack the worms. Worm lysates were boiled with 10μl 2x ample buffer at 95℃ for 15 minutes. Samples were separated with 15% SDS-PAGE and transferred to PVDF membrane (Bio-Rad, 1620177). After blocking with 5% milk in TBST for one hour under room temperature, membranes were incubated with primary antibodies diluted with 3% milk in TBST at 4 ℃ overnight. Then, the membranes were incubated with HRP-conjugated secondary antibody for 2 hours under room temperature and detected by Immobilon Western Chemiluminescent HRP Substrate (Millipore, WBKLS0500).

The following antibodies were used for Western blot analysis: rabbit anti-SMZ (Gene Script) at 1:5000 dilutions, rabbit anti-α-tubulin (Proteintech, 11224-1-AP) at 1:2000 dilutions, mouse anti-MSP (4A5, Hybridoma bank) at 1:500 dilutions, HRP-conjugated goat anti-mouse IgG (Millipore, AP124P) at 1:5000 dilutions, and HRP-conjugated goat anti-rabbit IgG (Millipore, AP132P) at 1:5000 dilutions. Western blot imaging was recorded by Bio-Rad ChemiDoc XRS^+^ Image System.

#### Electron microscopy

Synchronized day1 adult male worms were immobilized with 2μM levamisole and picked on the grid (10∼15 animals per grid) with eyelash and mixed with fresh OP50. Then the samples were subjected to cryofixation immediately with a high-pressure freezer EM HPM100 (Leica). Followed by subsequent freeze substitution and low temperature embedding with AFS2 EM (Leica). Sections were placed on 0.4% Formvar-coated slot grids and imaged with 80kV on a TEM (FEI Tecnai G2 Spirit TWIN) through a digital CCD camera. The images were analyzed with ImageJ Fiji and were processed with Adobe Illustrator 2025 software.

## Results

### SMZ-1/2 are redundant male-specific fertility factors

SMZ-1 and SMZ-2 are highly similar proteins, each containing two PDZ domains (Figure S1,[31]). To determine whether they function redundantly in spermatogenesis, we compared brood size and embryo hatch rates between wild-type and *smz* mutants. Worms carrying single mutations in either *smz-1* or *smz-2* showed brood sizes comparable to wild type (Figure 1A) and generated embryos with normal hatch rates (Figure 1B). By contrast, *smz-1; smz-2* double mutant hermaphrodites were completely sterile and laid unfertilized oocytes. Thus, SMZ-1 and SMZ-2 are functionally redundant for fertility (Figures 1AB and S2B). Consistent with these observations, DIC microscopy revealed normally developing embryos in the uteri from wild type, *smz-1,* and *smz-2* hermaphrodites (Figure 1C), whereas *smz-1; smz-2* hermaphrodites contained only unfertilized oocytes (Figure 1C). The sterility in *smz-1; smz-2* hermaphrodites was paternally originated, as mating with wild type males restored the brood size and embryonic hatch rates and resulted in normally developing embryos in the uterus (Figure 1 ABC). Given their redundant functions, we refer to SMZ-1 and SMZ-2 collectively as SMZ-1/2.

**Figure 1.**
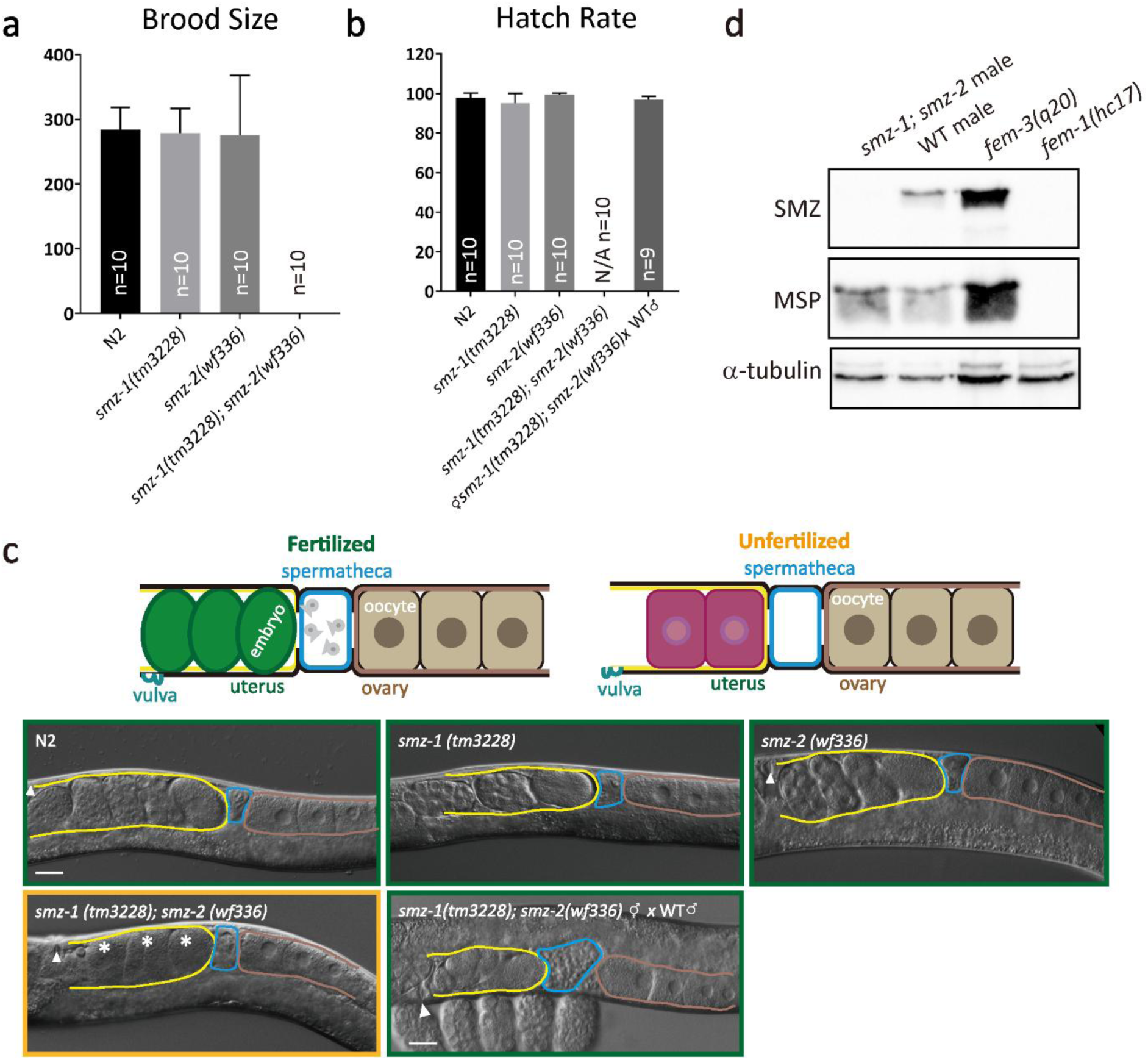
SMZ-1/2 are functionally redundant male-fertility factors. (A) Average number of brood sizes produced by N2, *smz-1(tm3228)*, *smz-2(wf336)*, and *smz-1(tm3228); smz-2(wf336)* hermaphrodites from the L4 stage through adulthood. *n* = 10 animals per group. Data are presented as mean ± SD. (B) Embryo hatch rates from the same genotypes as in (A), either unmated or mated with wild-type males for 16 hours, followed by egg laying over a 48-hour period. Data are presented as mean ± SD. (C) Top: schematic illustrations of hermaphrodites showing fertilized versus unfertilized reproductive tracts. Bottom: representative DIC images of N2, *smz-1(tm3228)*, *smz-2(wf336)*, and *smz-1(tm3228); smz-2(wf336)* hermaphrodites, either unmated or mated with wild-type males. *n* = 10 animals per group. Ovary, spermatheca, and uterus are outlined in brown, blue, and yellow, respectively. Arrowheads indicate the vulva; stars denote unfertilized oocytes in the uterus. (D) Western blot analysis of SMZ-1/2 protein levels in *smz-1/2* males, wild-type males, *fem-1(hc17)* hermaphrodites, *fem-3(q20)* hermaphrodites. MSP served as a male gonad marker; tubulin was used as a loading control. Each sample contained 100 hand-picked day1 adult animals.

To confirm that SMZ-1/2 function in spermatogenic germ cells, we examined their expression in spermatogenic versus oogenic germlines using an antibody against the shared C-terminal region [31]. SMZ-1/2 proteins were detected exclusively in sperm-producing *fem-3(q20)* extracts, with no detectable expression in oogenic germ cells or somatic tissues (Figure 1D). Together, these results establish SMZ-1 and SMZ-2 as male-specific fertility factors essential for functional sperm production.

### *smz-1/2* mutants arrested as primary spermatocytes

To further investigate the roles of SMZ-1/2 in male fertility, we compared DAPI-stained wild-type and *smz-1/2* male gonads and examine the developing spermatogenic nuclei. As previously documented, the post-pachytene wild-type spermatogenic gonad exhibited an array of nuclei with distinct morphological features representing diplotene, karyosome, dividing chromosomes, and mature sperm [37]. (Figure 2A). In stark contrast, *smz-1/2* mutant gonads showed an accumulation of abnormal nuclei in the division zone and lack of mature sperm (Figure 2A), indicating defects in male meiotic divisions. To determine whether these nuclear defects in *smz-1/2* mutants corresponded to meiotic arrest, we examined the dividing spermatocytes in isolated male gonads (Figure 2B). DIC microscopy revealed that dissected wild-type male gonads host a series of spermatocytes at primary or secondary dividing stages as well as mature sperm. In *smz-1/2* gonads, however, only primary spermatocytes with undivided nuclei were observed, strongly indicating a failure to complete meiotic division. These findings indicate that SMZ-1/2 are required for meiotic division progression during spermatocyte development.

**Figure 2.**
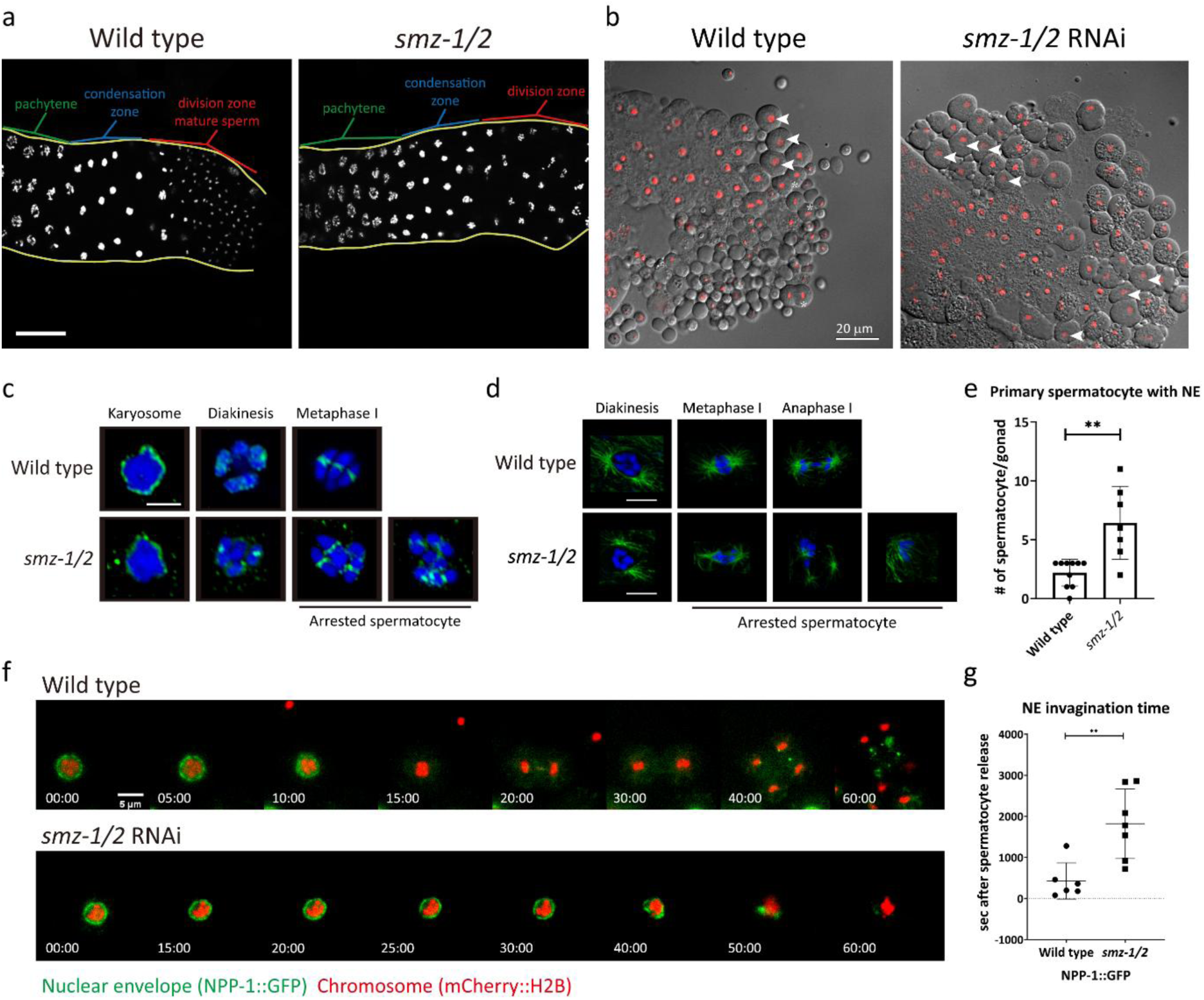
*smz-1/2* results in arrested primary spermatocytes. (A) DAPI-stained male gonads from wild-type and *smz-1/2* animals. Gonad boundaries are outlined in yellow. Spermatocyte developmental stages, pachytene, condensation zone, and division zone, are indicated in green, blue, and red, respectively. Scale bar: 20 μm. (B) Representative DIC images of released male gonads from wild-type and *smz-1/2* RNAi-treated animals. Nuclei were labeled with mCherry::H2B. Primary spermatocytes are marked with white arrowheads. Scale bar: 20 μm. (C) Individual developing spermatocytes at karyosome to metaphase I stages in wild-type and *smz-1/2* animals, stained with pH3S10 (green). Nuclei were counterstained with DAPI (blue). Scale bars: 5 μm. (D) Dividing spermatocytes at diakinesis to anaphase I stages in wild-type and *smz-1/2* animals, stained with α-tubulin (green) and DAPI (blue). Scale bars: 5 μm. (E) Quantification of primary spermatocytes retaining the nuclear envelope after release from wild-type and *smz-1/2* male gonads. Wild-type: *n* = 10. *smz-1/2*: *n* = 7. Data are presented as mean ± SD. \*\**P*<0.01. (F) Time-lapse imaging of primary spermatocytes after release from male gonads in wild-type and *smz-1/2*. Nuclear envelope (NE) was visualized using NPP-1::GFP, and nuclei were labeled with mCherry::H2B. Wild type: *n* = 6. *smz-1/2*: *n* = 7. (G) Quantification of NE invagination time in primary spermatocytes after gonad release. Data are presented as mean ± SD. \*\**P*<0.01.

### SMZ-1/2 are required for timely NEBD and meiotic progression

To determine whether SMZ-1/2 are essential for M phase progression, we examined the distribution of M phase marker phospho-histone H3 (pH3S10) in wild-type and *smz-1/2* male gonad by immunofluorescence staining [37, 38]. In wild-type, pH3S10 localized to karyosome-stage nuclei and later to metaphase I bivalents (Figure 2C). Despite their meiotic arrest, *smz-1/2* spermatocytes displayed similar pH3S10 localization patterns, indicating successful M phase entry (Figure 2C). Furthermore, as wild-type spermatocytes assemble bipolar microtubule spindles, comparable structures were observed in diakinesis-stage *smz-1/2* spermatocytes (Figure 2D, [37]). Interestingly, arrested *smz-1/2* primary spermatocytes often showed aberrant spindle morphologies, including multipolar and disorganized microtubule arrays (Figure 2D, Movie S1&S2). These results suggest that meiotic spindles were assembled as *smz-1/2* spermatocytes progressed into M phase but could not be maintained, leading to failure of chromosome segregation.

While analyzing DIC images, we observed that most of *smz-1/2* primary spermatocytes retained their nuclear envelopes more frequently than wild-type (Figure 2BE), indicating defective nuclear envelope breakdown (NEBD). To test this, we performed time-lapse imaging using the NE marker NPP-1::GFP (Figure 2F). In wild-type, NEBD occurred rapidly, with NPP-1::GFP dispersing into cytoplasmic foci, followed by chromosome segregation and sperm formation. In contrast, *smz-1/2* spermatocytes showed significantly delayed NEBD and abnormal NPP-1 dispersal with failure of chromosome segregation initiation (Figure 2FG, Movie S3&S4). These results suggest that SMZ-1/2 act upstream of NEBD to ensure timely disassembly of the nuclear envelope during spermatocyte maturation.

### SMZ-1/2 localize as cytoplasmic foci from diplotene through spermatid formation

Given the division failures and delayed NEBD in *smz-1/2* mutants, we asked whether SMZ-1/2 localize to the nuclear envelope or other division-related structures. Immunofluorescence staining with antibodies recognizing the C-terminus of SMZ-1/2 revealed that SMZ-1/2 were first detected in the late syncytial spermatogenic germline, spanning diplotene to mature sperm stages. (Figure 3A). This pattern was confirmed by FLAG immunostaining of 3×FLAG::SMZ-1 males (Figure 3B). High-magnification imaging showed that SMZ-1/2 form dynamic cytoplasmic foci, initially as small puncta in diplotene, which elongate into bar-like structures, and later dispersed into cytoplasm in mature sperm. (Figure3C). To further investigate the dynamics of SMZ-1/2 foci during late spermatogenic stages, we constructed whole-cell 3D SMZ-1/2 structures and quantified their volume and number from diplotene to primary spermatocyte stages (Figure 3DE). The volume and number of SMZ-1/2 foci increased as spermatocytes progressed toward division. Together, these results suggest that SMZ-1/2 undergo cytoplasmic reorganization during spermatocyte maturation, possibly coordinating meiotic progression and cytoplasmic changes.

**Figure 3.**
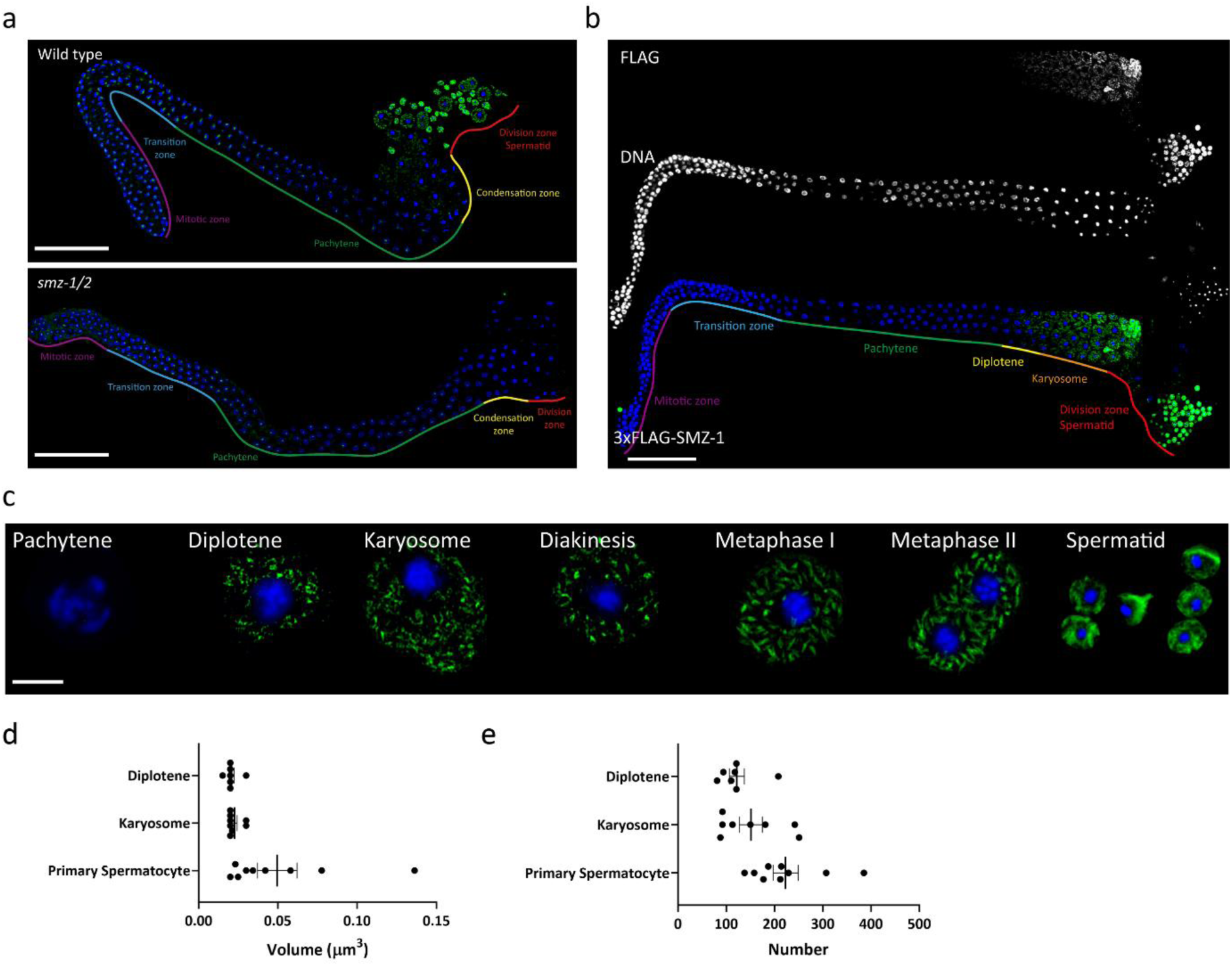
SMZ-1/2 distribution and subcellular localization in the male germline. (A) Released male gonads co-labeled with anti-SMZ (green) and DAPI (blue) in both wild-type and *smz-1/2* males. The developmental stages are labeled with indicated colors. Comparing between the wild-type and *smz-1/2* gonads, SMZ-1/2 signals distributed from condensation zone to mature sperm. Scale bar: 50 μm. (B) Released male gonads from 3xFLAG::SMZ-1 males co-stained with anti-FLAG (green) and DAPI (blue). The results are consistent with (A). Scale bar: 50 μm. (C) Subcellular localization of SMZ-1/2 during spermatocyte development from isolated wild-type spermatocytes and spermatids. The cells were labeled with SMZ (green) and DAPI (blue). Scale bar: 5 μm. (D) Measurements of SMZ-1/2 average volume in wild-type from diplotene to primary spermatocytes. *n* = 7 cells in diplotene. *n* = 8 cells in karyosome. *n* = 9 cells in primary spermatocyte (includes diakinesis to metaphase I cells). (E) Average SMZ-1/2 particle numbers in wild-type from the stages in (D).

Results from SMZ-1/2 localization analyses argued against our initial hypothesis that SMZ-1/2 might localize to the nuclear envelope (NE) to facilitate timely NEBD. Co-immunostaining with nucleoporins in primary spermatocytes preparing for NEBD revealed that SMZ-1/2 signals were not detected at the NE (Figure S3AB). These findings suggest that SMZ-1/2 likely regulate NEBD indirectly through cytoplasmic functions.

### SMZ-1/2 are associated with MSP during FB-MO development

While examining the subcellular localization of SMZ-1/2, we noticed that these proteins translocated from the cytoplasm to the pseudopods during sperm activation, similar to MSP, the main cytoskeletal component for the ameboid sperm movement [2, 35]. This resemblance led us to hypothesize that SMZ-1/2 are novel components of the FB-MO and may associate with this structure to regulate spermatogenic processes. To test this, we co-labeled SMZ-1/2 and FB-MO and analyzed their distribution across spermatogenic developmental stages (Figure 4A). Whole-cell level analysis revealed that SMZ-1/2 signals were broadly coincident with FB-MO localization from diplotene to diakinesis. Colocalization analysis revealed that SMZ-1/2 were initially associated with both FB and MO structures during diplotene, but their correlation with the MO decreased as cells progressed through karyosome and diakinesis (Figure 4B). In contrast, SMZ-1/2 maintained a consistently strong correlation with MSP filaments throughout these stages.

**Figure 4.**
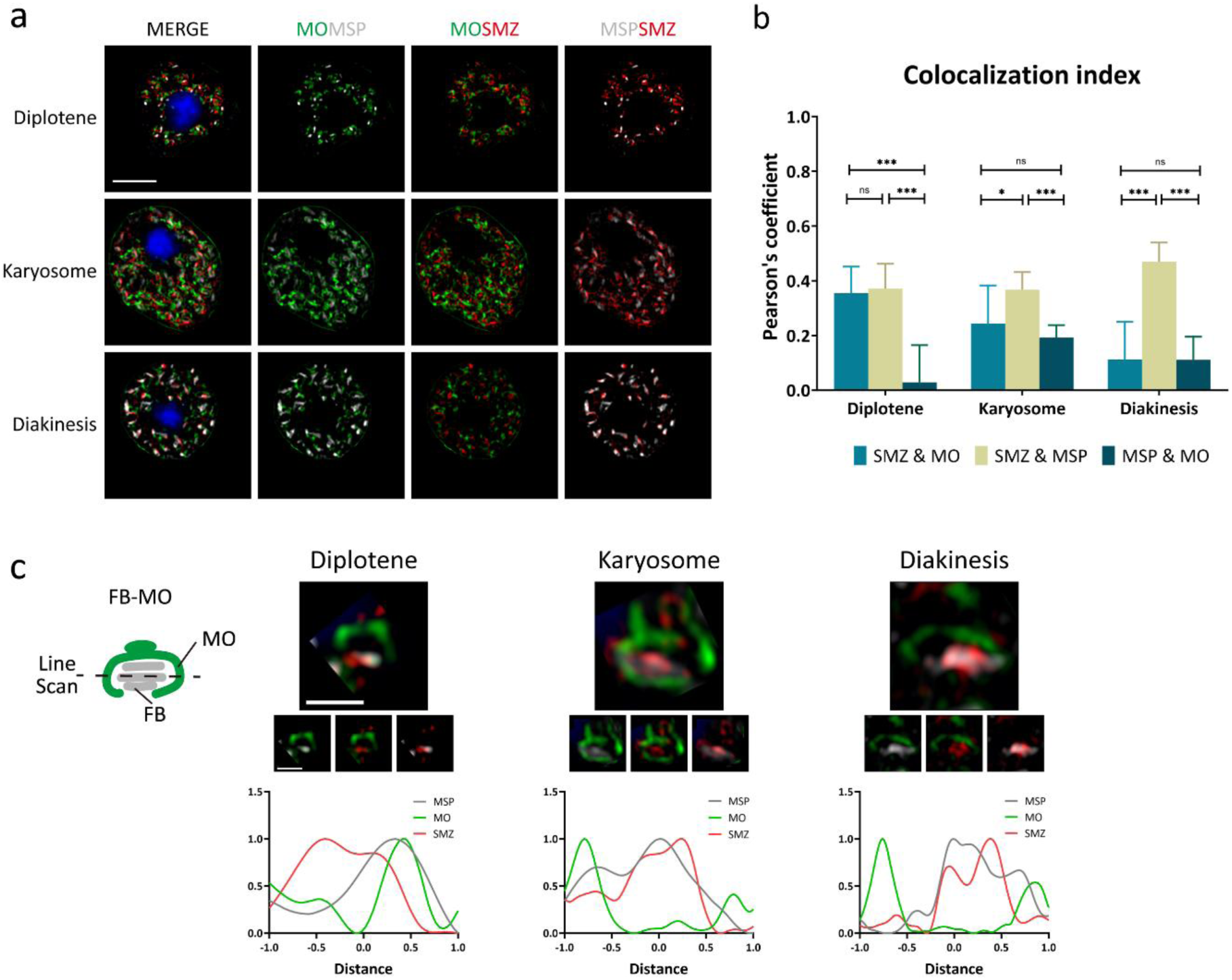
SMZ-1/2 are associated with FB during FB-MO development. (A) Isolated developing spermatocytes from wild-type males at diplotene to diakinesis stages, co-stained with SMZ (red), MO (green), MSP (gray), and DAPI (blue). Scale bar: 5 μm. (B) Quantification of colocalization index between SMZ and FB-MO signals from (A). Diplotene: *n* = 9. Karyosome: *n* = 11. Diakinesis: *n*=11. Data are presented as mean ± SD. \*\*\**P* < 0.001; \**P* < 0.05; ns, *P* > 0.05. (C) Left: Schematic illustration of FB-MO. Upper right panel: Representative single FB-MO shown in (A). The small boxes in the middle represent: MSP/MO, SMZ/MO, and SMZ/MSP from left to right. Scale bar: 1 μm. Bottom right: Corresponding fluorescence intensity line scan profile of each individual FB-MO.

We next inspected the spatial relationship between SMZ-1/2 and FB-MO in individual FB-MOs across developmental stages and performed fluorescence intensity profiling aligned with SMZ-1/2 (Figure 4CD). As previously described [17], MSP filaments appeared as small cytoplasmic foci during the diplotene stage and elongated into characteristic bar-like structure during karyosome and diakinesis (Figure 4C). Notably, SMZ-1/2 signals closely tracked this elongation, forming parallel structures that mirrored MSP filament growth. Line scan analyses revealed that SMZ-1/2 did not preferentially associate with either the MO or MSP in the diplotene stage. However, in the karyosome and diakinesis stages, SMZ-1/2 became more tightly associated with MSP filaments within the MO compartment. Together, these findings indicate that SMZ-1/2 dynamically associate with MSP filaments during FB-MO development and likely contribute to MSP filament assembly in developing spermatocytes.

### *smz-1/2* failed to form MSP filaments during late spermatogenesis

Because SMZ-1/2 are associated with MSP, we asked whether these proteins are essential for MSP filament development. We first examined FB-MO distribution in wild-type and *smz-1/2* male gonads (Figure 5A). In wild-type, FB-MOs are distributed from syncytium to the mature sperm. In *smz-1/2*, however, the FBs were absent despite the presence of MOs emerging from the syncytium. We next examined the FB-MO structures in wild-type and *smz-1/2* spermatocytes and performed whole-cell 3D reconstructions to quantify MSP-containing structures (Figure 5B). In wild-type spermatocytes, FB-MO volume increased as cells progressed from diplotene to diakinesis, consistent with the pattern observed in individual FB-MOs (Figure 4C). MSP volume significantly expanded as cells reached the karyosome stage (Figure 5C), whereas the number of MSP structures remained relatively constant (Figure 5D), suggesting that filament quantity is established early, while filament elongation is dynamically regulated. In contrast, MSP filaments were absent in *smz-1/2* spermatocytes, and the volume of MO did not change across stages (Figure 5B), indicating that MSP filament assembly was disrupted despite the presence of MSP protein (Figure 1D). These findings suggest that SMZ-1/2 are required for initiating MSP filament assembly.

**Figure 5.**
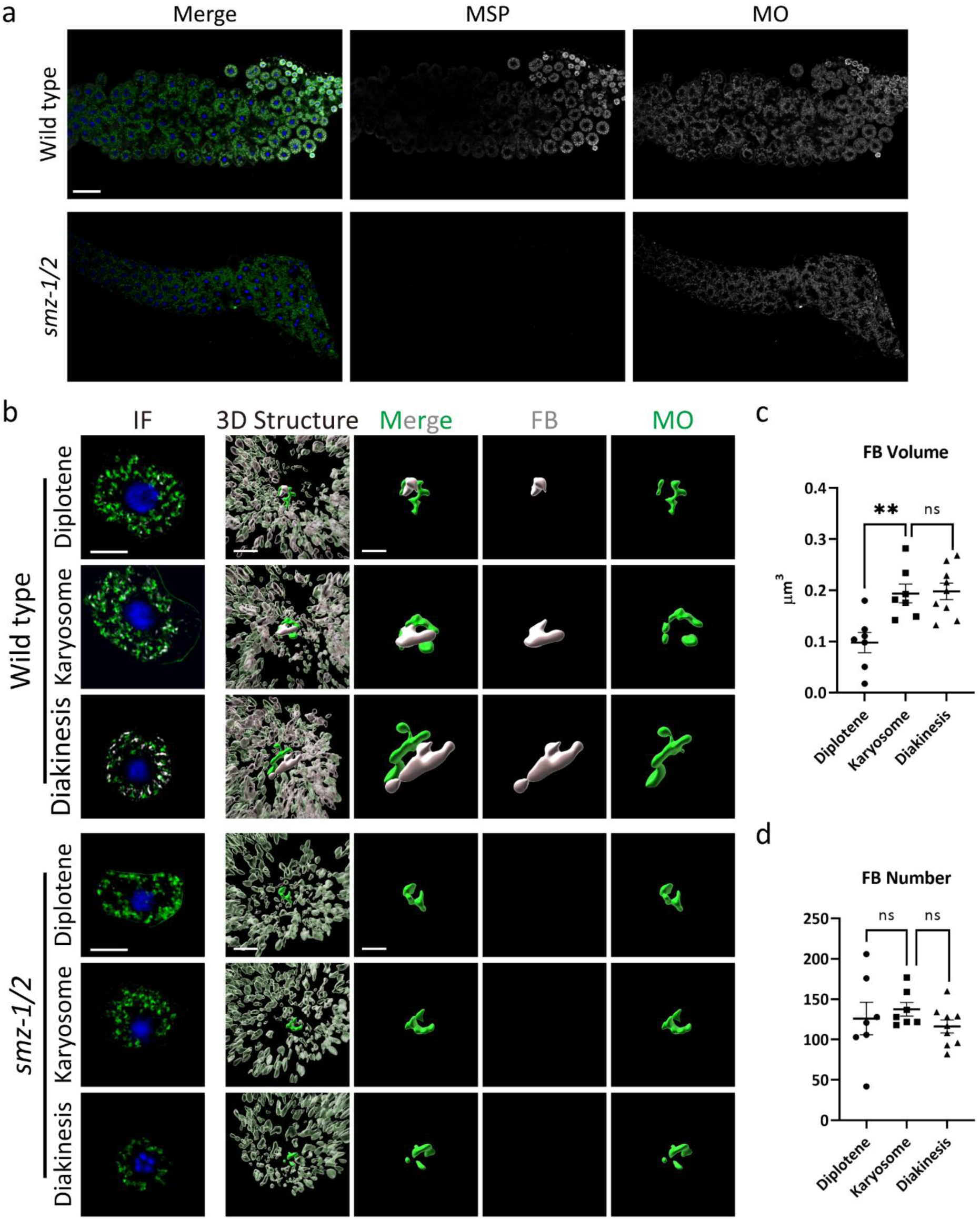
SMZ-1/2 are essential for FB-MO development. (A) Gonads from wild-type and *smz-1/2* co-labeled with MO (green), MSP (gray), and DAPI (blue). Scale bar: 20 μm. (B) Isolated developing spermatocytes filmed with confocal microscopy to identify the subcellular morphology of FB-MOs from wild-type and *smz-1/2* males at diplotene to diakinesis stages. Scale bar: 5 μm. The 3D reconstructions of both wild-type and *smz-1/2* FB-MOs from images in the left IF panel. FB-MO of interest is shown in solid color, while surrounding FB-MOs are rendered with increased transparency. Scale bar: 2 μm. The right three panels are single FB-MOs from the construction images. Scale bar: 1 μm. (C) (D) FB size measurement and number quantification per cell based on 3D reconstructions from wild-type spermatocytes in (A). (C) Average FB size from each stage. (D) Total FB number. *n* = 7 cells in both diplotene and karyosome groups. *n* = 9 cells in diakinesis group. Data are presented as mean ± SEM. \*\**P* < 0.01; ns, *P* > 0.05.

### *smz-1/2* mutants display and smaller MOs during FB-MO development

To further investigate how SMZ-1/2 affect FB-MO development, we examined FB-MO structures in wild-type and *smz-1/2* male germlines using transmission electron microscopy (TEM). In wild-type spermatocytes, MOs first appeared in pachytene as double-membraned structures, then progressively filled with crystalline FBs in diplotene and elongated further in dividing spermatocytes before shrinking again in spermatids (Figure 6A). These results were consistent with our immunofluorescence analyses that MSP progressively elongates inside the MO (Figure 4AC, [17]).

**Figure 6.**
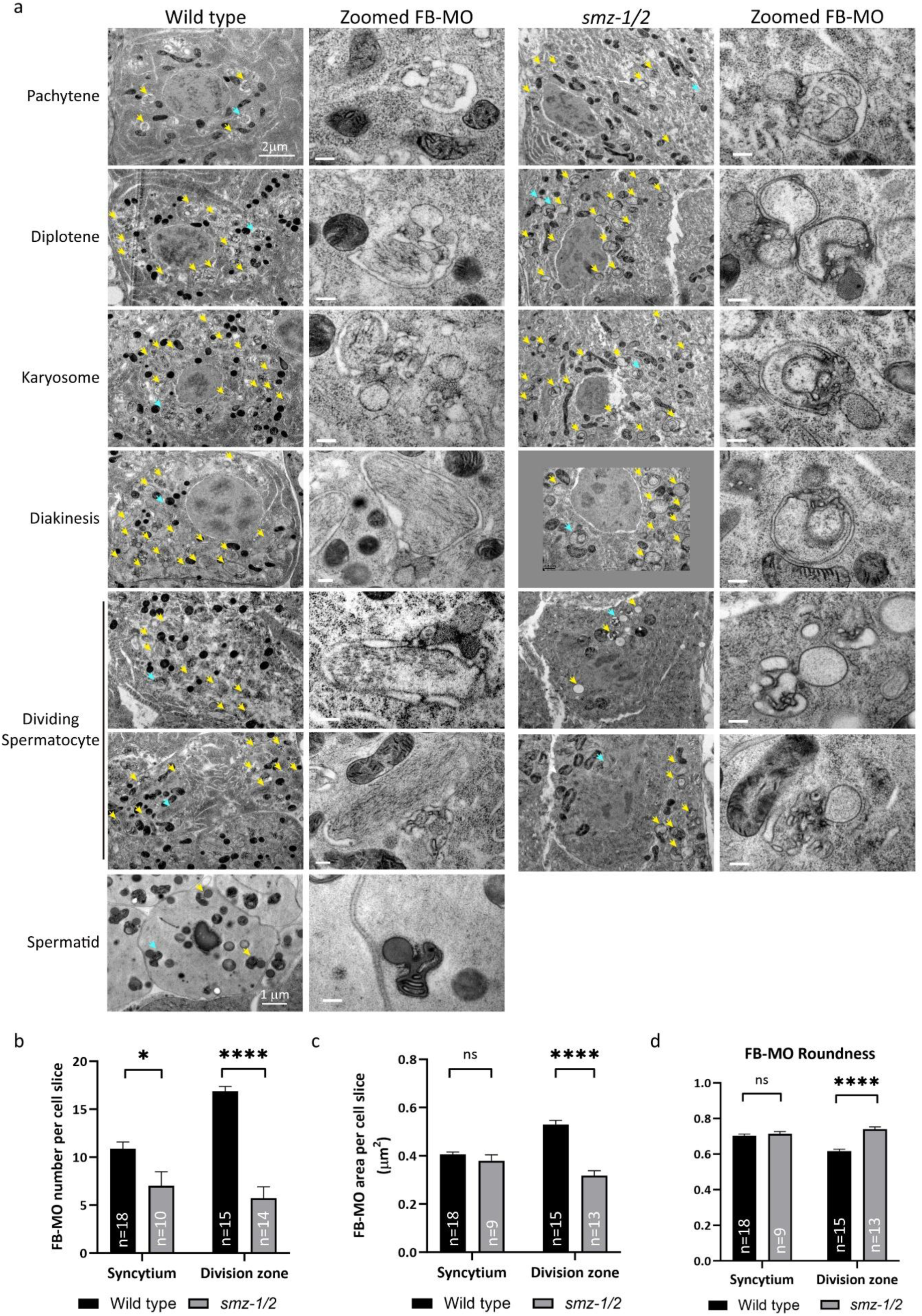
*smz-1/2* spermatocytes lack MSP filaments. (A) Transmission electron micrographs (TEM) of selected stages of sperm development in wild-type and *smz-1/2* animals. For each group, the left panels show entire spermatocytes, and the right panels display both enlarged and isolated FB-MOs. Scale bars: 2 μm for whole-cell images (unless otherwise indicated); 200 nm for the isolated single FB-MOs. (B) (C) (D) FB-MO quantification per EM slice in both wild-type and *smz-1/2* from (A). Cells were categorized as belonging to either the syncytial region or division zone. (B) Total FB-MO numbers per EM slice. (C) Average FB-MO area. (D) FB-MO roundness analyses. Data are presented as mean ± SEM. \*\*\*\**P* < 0.0001; \**P* < 0.05; ns, *P* > 0.05.

In *smz-1/2* mutants, MOs also formed as double-membrane structures in pachytene, indicating that initial membrane formation took place. However, crystalline FBs were never observed inside MOs at later stages. Instead, MO bodies remained empty, failed to elongate, and appeared underdeveloped by the division stage (Figure 6A). Quantification showed that *smz-1/2* mutants produced significantly fewer FB-MOs than wild-type, and those present were smaller and more rounded than in wild-type (Figure 6CD). These results show that SMZ-1/2 are primarily required for MSP filament incorporation into MOs. The immature MO morphology in *smz-1/2* mutants likely reflects MO atrophy as a result from failure of MSP filament loading.

### SMZ-1/2 function downstream of casein kinase SPE-6 to promote MSP filament formation

SPE-6 is a male-specific casein kinase that is essential for MSP filament assembly [23]. Because *smz-1/2* mutants exhibited phenotypes similar to *spe-6* mutants, we hypothesized that SMZ-1/2 might act downstream of SPE-6 to regulate MSP filament formation. To test this, we examined the subcellular localization of SMZ-1/2 in *spe-6* mutant males (Figure 7). We found that, in contrast to the distinct bar-like cytoplasmic structures observed in wild-type, SMZ-1/2 were diffusely distributed throughout the cytoplasm in *spe-6* mutants (Figure 7). These results show that SPE-6 is required for proper SMZ-1/2 localization. To further determine whether SMZ-1/2 and SPE-6 also influence MO organization, we analyzed FB-MO distribution and subcellular localization (Figure 8). In *smz-1/2*, *spe-6*, and *smz-1/2; spe-6* males, the distribution of MOs were comparable to wild-type, while MSP signals were completely absent. These results indicate that SPE-6 is necessary for SMZ-1/2 cytoplasmic structures, which are in turn required for MSP filament initiation.

**Figure 7.**
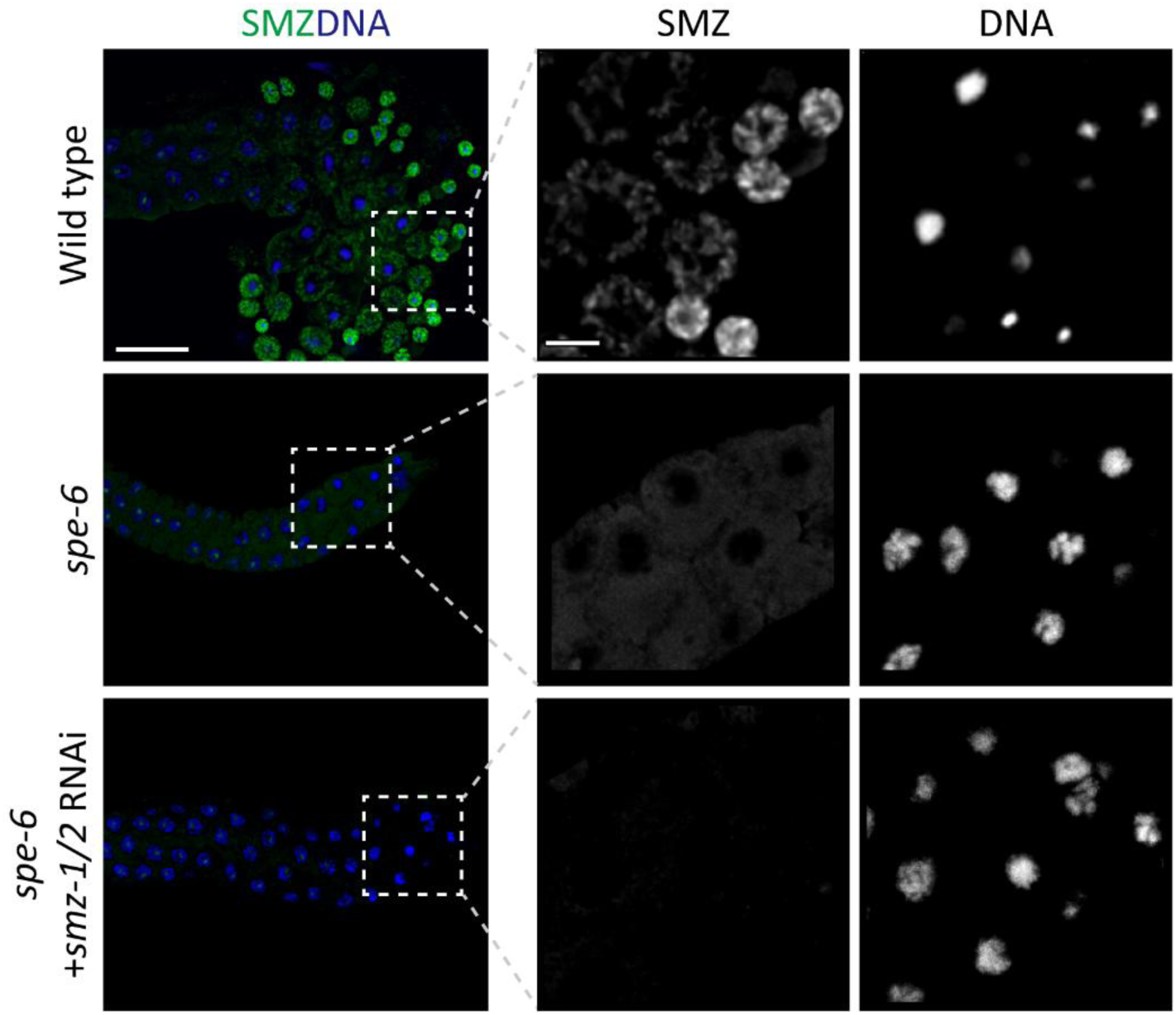
SPE-6 is required for SMZ-1/2 subcellular structure. Isolated male gonads that stained with SMZ-1/2 and DNA in both wild type and *smz-1/2*. The zoomed area is circled with white dash line. The wild-type showed SMZ-1/2 cytoplasm distribution. *spe-6* had SMZ signals dispersed in the cytoplasm. For the cropped gonad to the left scale bar is 20 μm, and the zoomed images are scaled in 5 μm.

**Figure 8.**
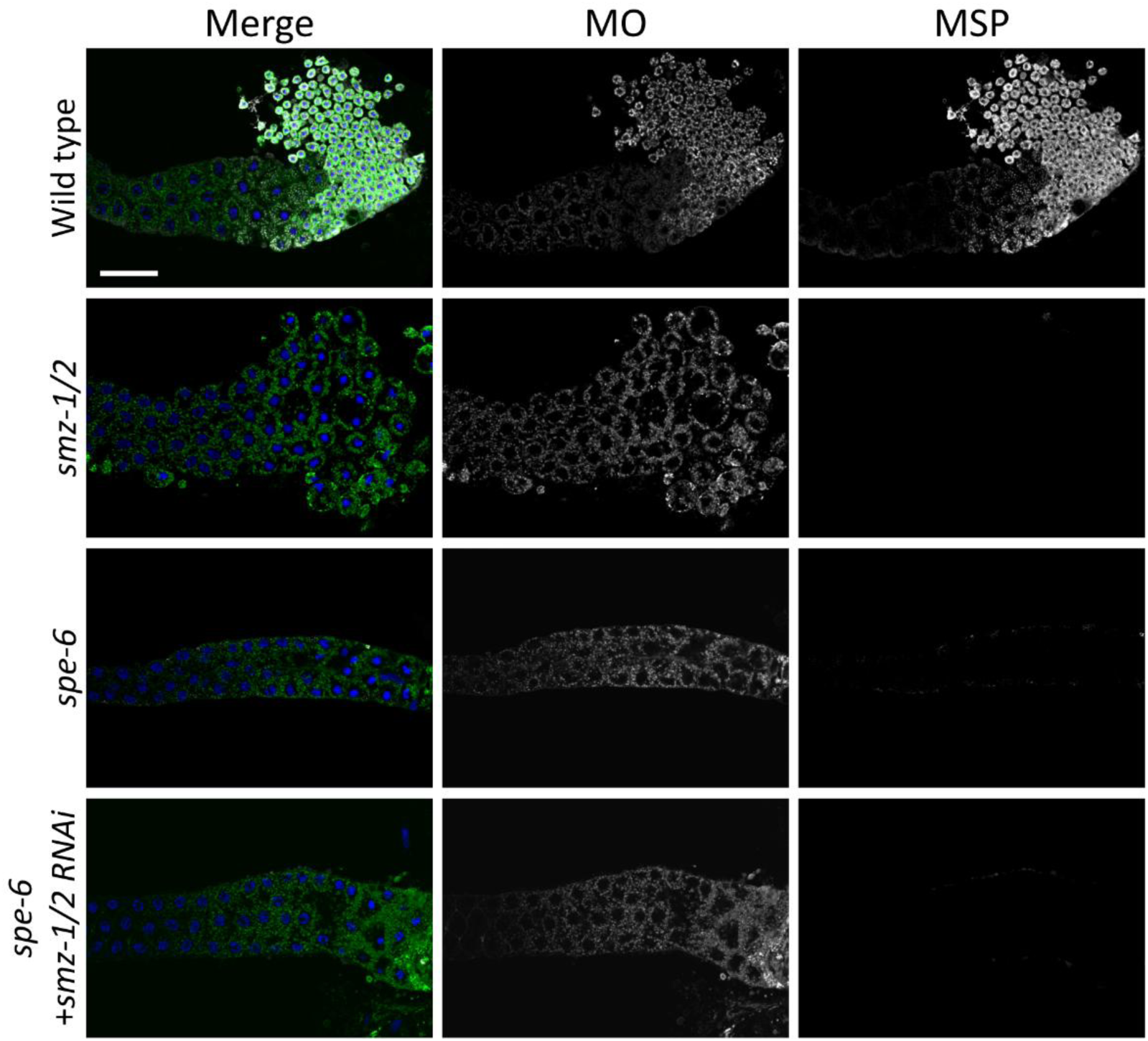
SPE-6/SMZ-1/2 pathway is independent from MO distribution. Immunolabeling of wild-type, *smz-1/2*, *spe-6*, and *spe-6* with *smz-1/2* depletion male gonads. The gonads were marked with MSP (grey), MO (green), and DAPI (blue). MO signals comparably distributed in the worms either with wild-type genome or with variant mutations. Scale bar: 20 μm.

## Discussion

In this study, we identify SMZ-1 and SMZ-2 as sperm-specific proteins that are required for the proper assembly of the fibrous body–membranous organelle (FB-MO), a compartment unique to nematode spermatogenesis. Our results show that SMZ-1/2, two PDZ domain–containing proteins expressed specifically in spermatogenic germline, associate with major sperm protein (MSP) and are necessary for the onset of MSP filament formation. In the absence of SMZ-1/2, MSP fails to assemble, FB-MOs are reduced and abnormal, and spermatocytes arrest before completion of meiosis. These phenotypes are likely regulated by SPE-6-depedent SMZ-1/2 structures. Taken together, these findings suggest that SMZ-1/2 act as key initiators of FB-MO formation and expand the growing set of factors that regulate MSP-dependent sperm development.

The meiotic defects observed in *smz-1/2* mutants underscore the importance of proper FB-MO formation in germline development. *smz-1/2* spermatocytes enter M phase and assemble spindles, but exhibit delayed nuclear envelope breakdown, fail to segregate chromosomes, and do not produce mature sperm. Similar primary spermatocyte arrest phenotypes are seen in other FB-MO mutants such as *spe-4* and *spe-6*, linking FB-MO integrity to meiotic progression [16, 23, 39]. Because FB-MOs are derived from the endoplasmic reticulum, these observations are consistent with broader evidence that endomembrane organization can influence chromosome segregation and nuclear division. Indeed, work in mitotic cells has shown that endoplasmic reticulum (ER) organization can influence chromosome segregation [40, 41], highlighting a broader connection between endomembrane dynamics and nuclear division. Future studies will be needed to define how FB-MO stability coordinates cytoskeletal regulation, endomembrane organization, and meiotic events during spermatogenesis.

Our data place SMZ-1/2 within the SPE-6-mediated MSP filament assembly network (Figure 9). The casein kinase SPE-6 is required for MSP filament initiation, as *spe-6* mutants retain MSP in the cytoplasm and lack filaments [23]. We find that SMZ-1/2 cytoplasmic structures are lost in *spe-6* mutants, indicating that SPE-6 activity is required for the formation or stability of SMZ-1/2 assemblies. In *Ascaris suum*, MSP polymerization is similarly promoted by kinase activity, where MAPK-associated MPOP phosphorylates MFP2 to stimulate filament growth [42]. Homologs of these factors, MSD-1 and NSPH-2, were recently identified in *C. elegans* and shown to colocalize with MSP filaments during spermatogenesis [43], highlighting a conserved role for kinase-mediated regulation across nematode spermatogenic stages. Within this framework, our data are consistent with the idea that SMZ-1/2 may act as potential scaffold proteins that facilitate the organization of MSP regulators at sites of FB-MO biogenesis. This idea is supported by our electron microscopy analyses showing that *smz-1/2* mutants lack MSP filaments and by the concomitant reduction in FB-MO number and aberrant FB-MO morphology, phenotypes consistent with defective recruitment of MSP into developing FB-MOs. Together, these findings support a defect at the initiation of MSP filament assembly step rather than a late polymerization failure.

**Figure 9.**
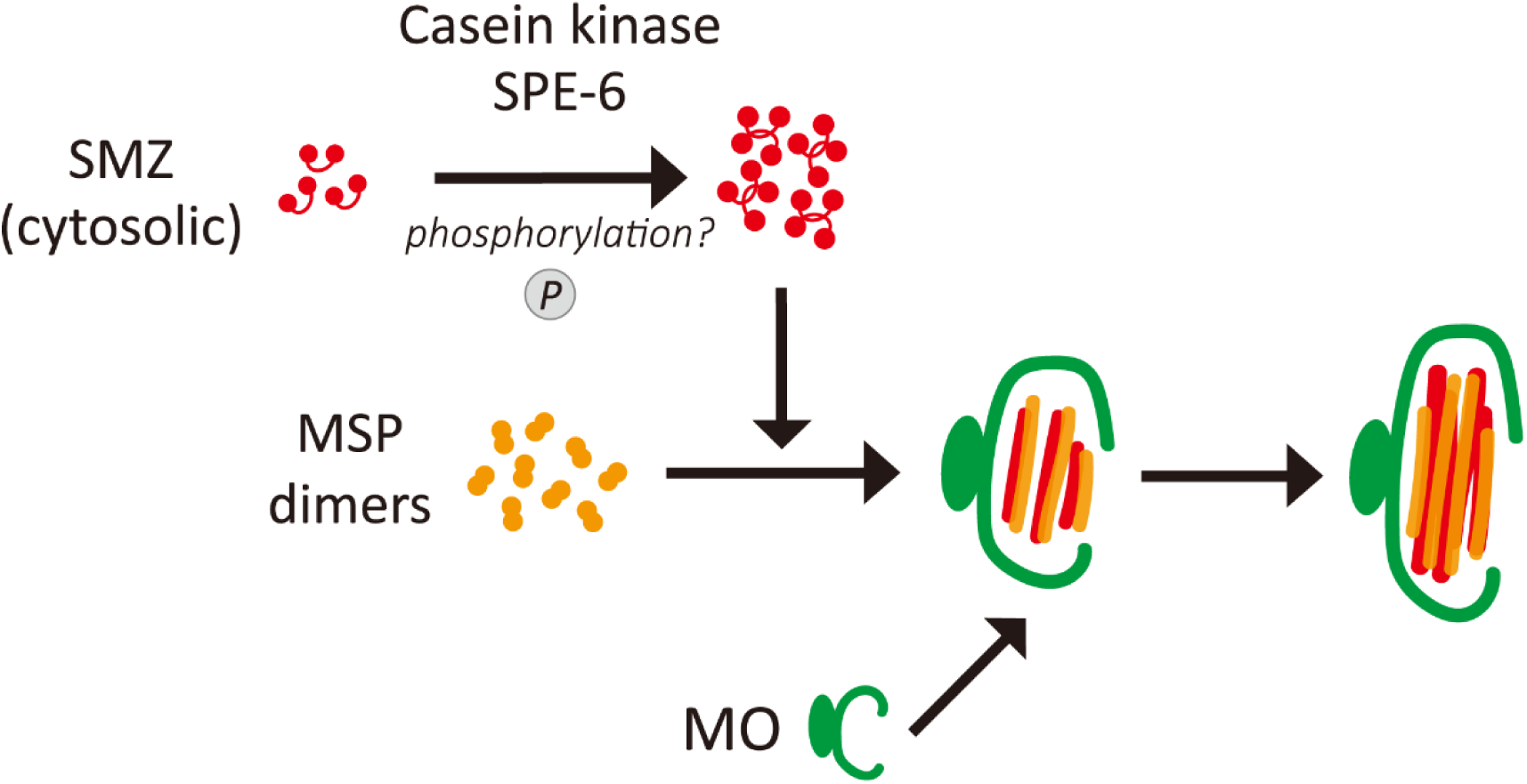
SMZ-1/2 in MSP filament assembly. SMZ-1/2 proteins act downstream of the casein kinase SPE-6 to form cytoplasmic structures. As SMZ-1/2 formed in the cytoplasm, they associated with MSP to initiate MSP filament assembly inside the MOs.

The predicted scaffolding role of SMZ-1/2 aligns with known functions of PDZ-domain proteins in organizing signaling and cytoskeletal complexes at membranes [24, 25, 44]. Across systems, PDZ scaffolds, such as PSD-95 and DLG/ZO family members, cluster receptors, kinases, and actin regulators in diverse contexts [45–47]. Furthermore, PDZ proteins have been implicated in acrosome reaction and junction remodeling during mammalian spermatogenesis [48–51]. We propose that SMZ-1/2 scaffold MSP and MSP regulators at FB–MO initiation sites, positioning kinase activity and other effectors to create a competence zone for MSP filament assembly. In this view, the empty, undeveloped MOs in *smz-1/2* mutants reflect a failure to load MSP filaments into the FB–MO compartment.

Together, our study identifies SMZ-1 and SMZ-2 as PDZ scaffold proteins that are required for the initiation of MSP filament assembly and FB-MO biogenesis in *C. elegans*. SMZ-1/2 expand the repertoire of MSP regulators and represent, to our knowledge, the first PDZ-containing candidates implicated in nematode sperm development. Future work to define SMZ-1/2 interaction partners and to test how disrupting PDZ binding or membrane targeting affects MSP loading will clarify how scaffolded signaling modules control organelle-based cytoskeletal assembly during spermatogenesis.

---

**Figure S1.**
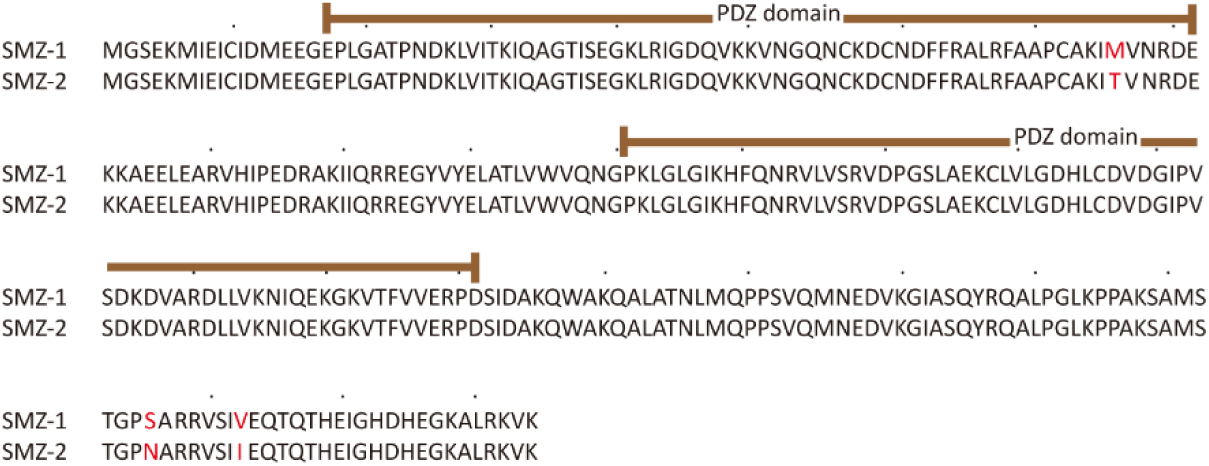
SMZ-1 and SMZ-2 share 99% sequence identity. Protein sequence alignment of SMZ-1 and SMZ-2. Brown lines indicate the positions of the PDZ domains. Red highlights denote amino acid differences between the two proteins.

**Figure S2.**
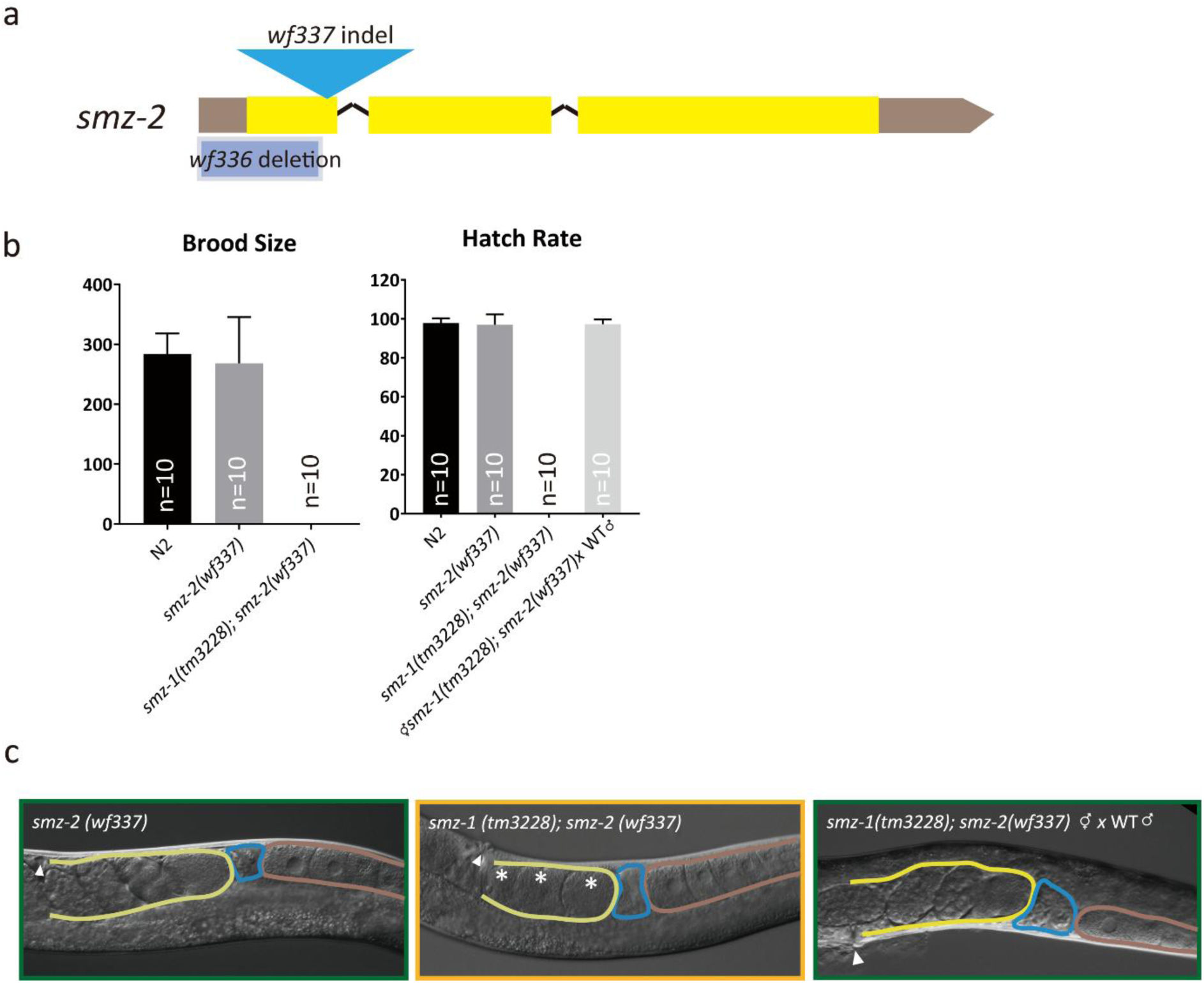
The alternative *smz-2* allele *wf337* exhibits a comparable phenotype to *wf336*. (A) Schematic illustration of the *smz-2* gene showing indels generated via CRISPR/Cas9. The *wf336* allele is a deletion that includes the start codon of *smz-2*, while the *wf337* allele is an indel that introduces a frameshift mutation. (B) Average brood sizes and embryo hatch rates of N2, *smz-2(wf337)*, and *smz-1(tm3228); smz-2(wf337)* hermaphrodites, either unmated or mated with wild-type males for 16 hours. Brood sizes were counted from L4 through adulthood. Hatch rates were calculated for eggs laid within 48 hours after mating. *n* = 10 animals per group. Data are presented as mean ± SD. (C) Representative DIC images of *smz-2(wf337)* and *smz-1(tm3228); smz-2(wf337)* hermaphrodites, either unmated or mated with wild-type males. *n* = 10 animals per group. Ovary, spermatheca, and uterus are outlined in brown, blue, and yellow, respectively. Arrowheads indicate the vulva; stars denote unfertilized oocytes in the uterus.

**Figure S3.**
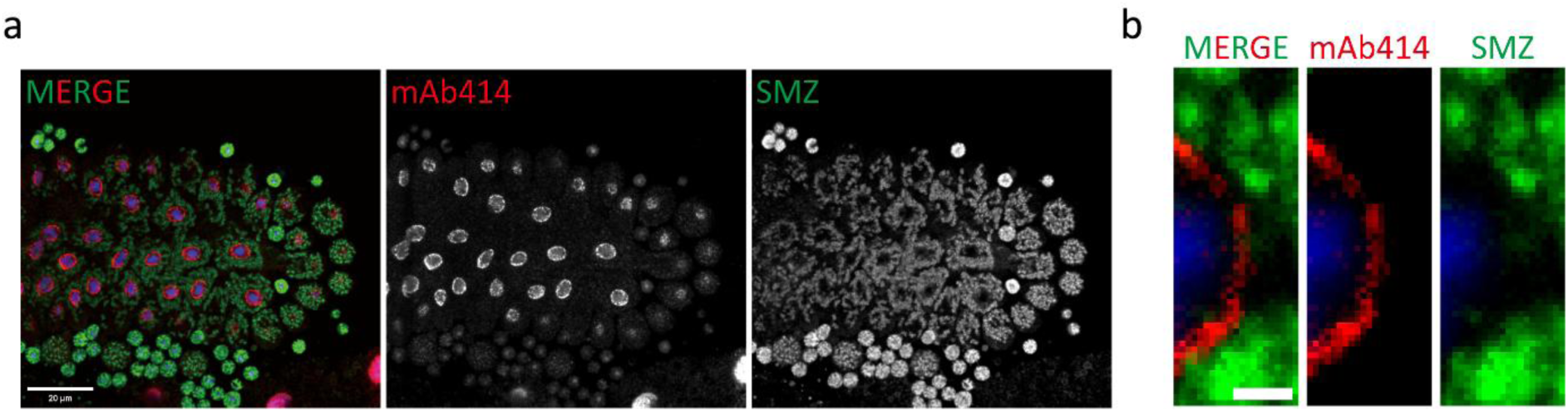
SMZ-1/2 do not localize to nuclear envelope. (B) Released male gonads co-labeled with anti-SMZ (green), mAb414 (red, nuclear envelope marker) and DAPI (blue) in wild-type males. Scale bar: 20 μm. (C) Magnified images of nuclear envelope from (A), showing the subcellular localization of both SMZ-1/2 and nuclear envelope. SMZ-1/2 do not localize to the nuclear envelope. Scale bar: 1 μm.

**Table S1:**
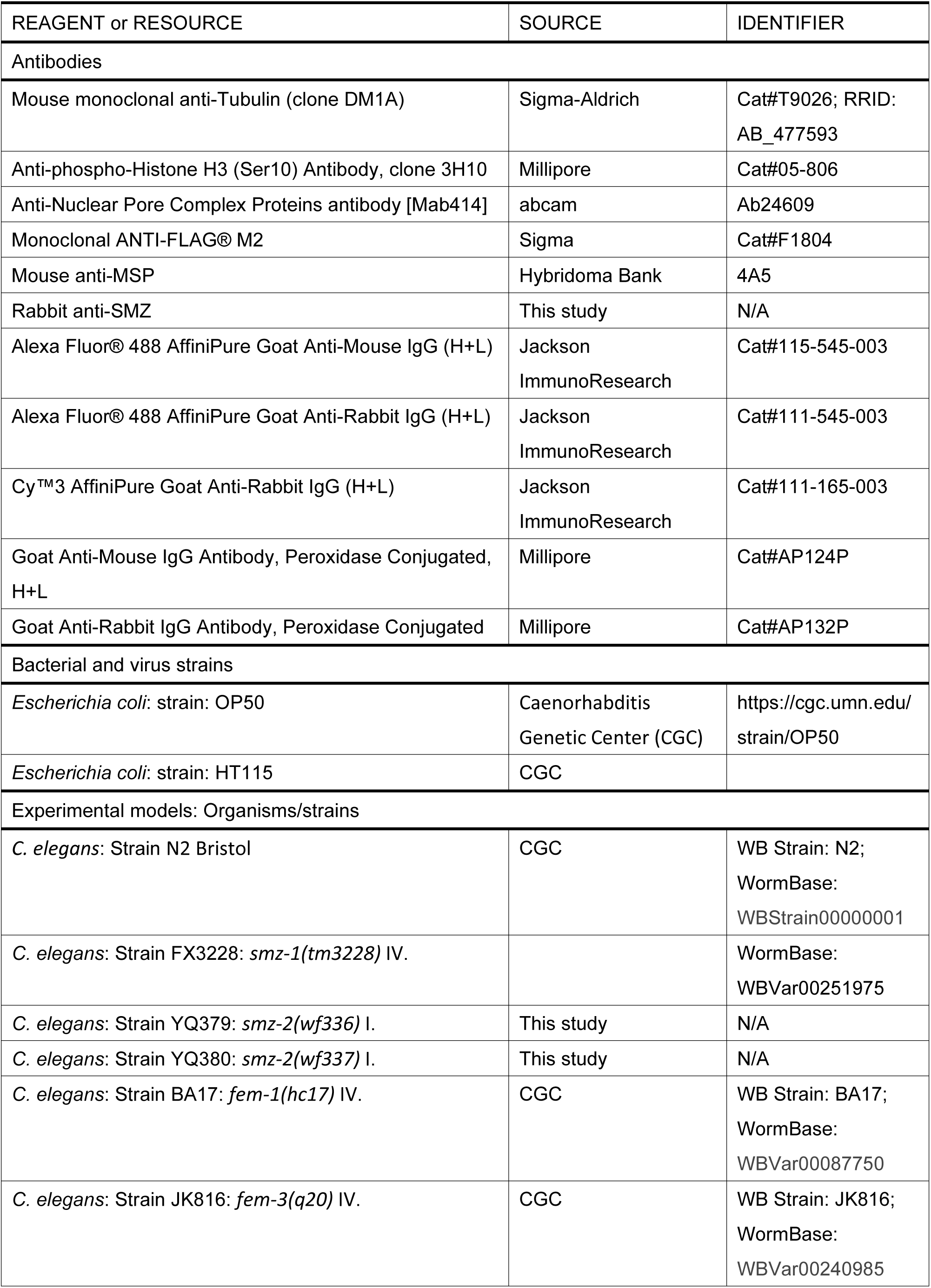
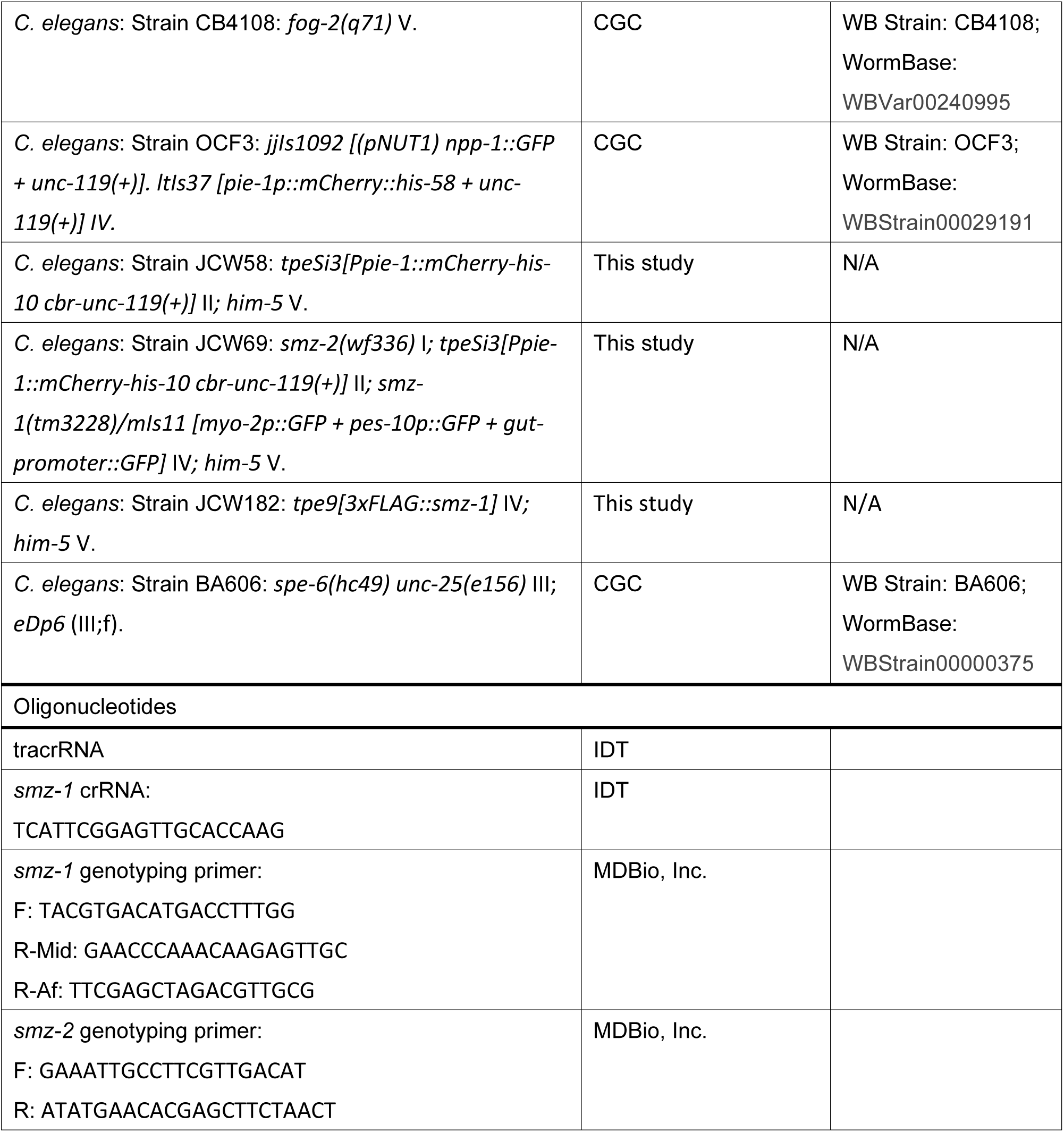
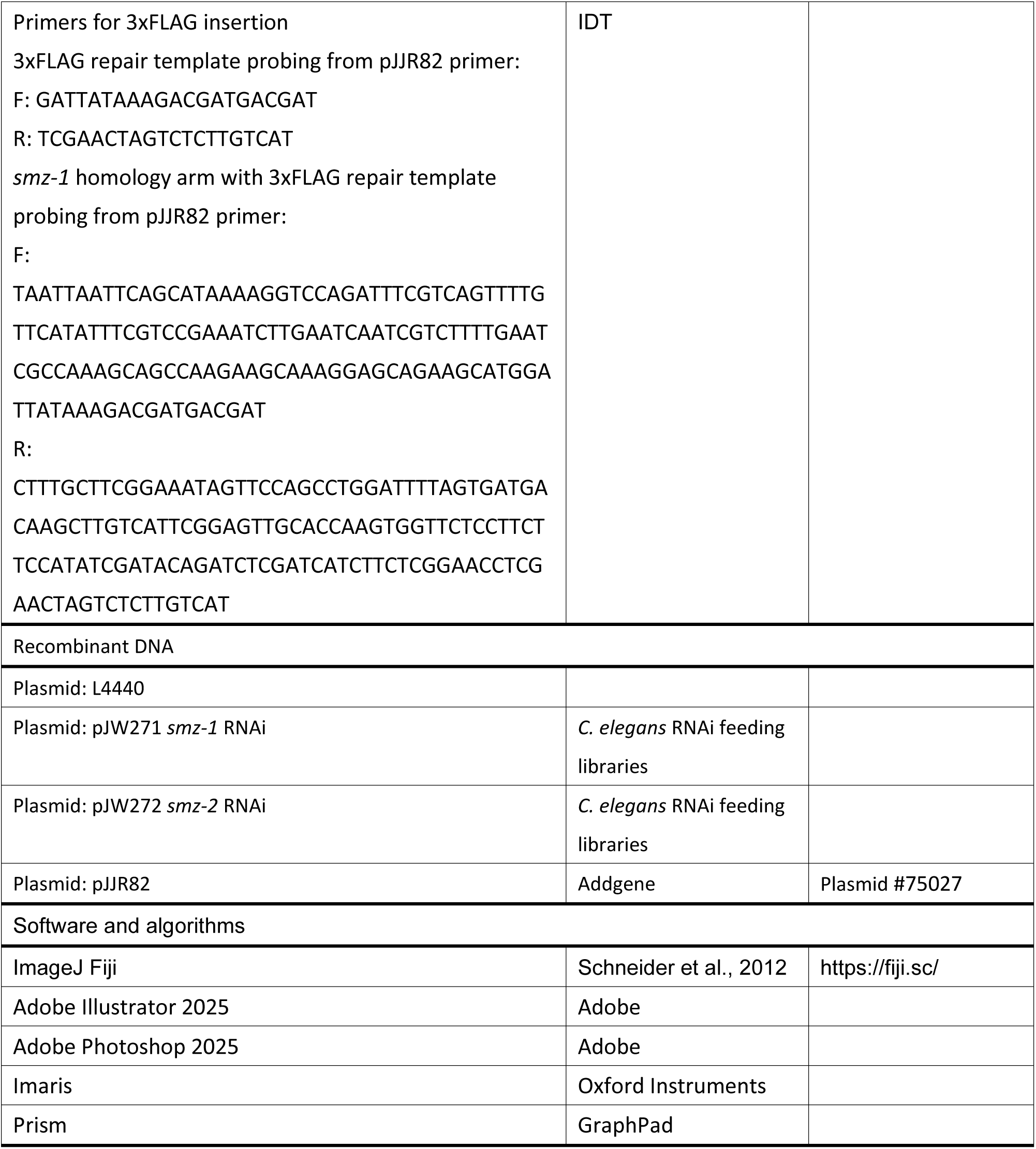
Key resources table.

## Reference

1. Gadadhar, S., T. Hirschmugl, and C. Janke, The tubulin code in mammalian sperm development and function. Semin Cell Dev Biol, 2023. 137: p. 26–37.

2. Sepsenwol, S., H. Ris, and T.M. Roberts, A unique cytoskeleton associated with crawling in the amoeboid sperm of the nematode, Ascaris suum. J Cell Biol, 1989. 108(1): p. 55–66.

3. Smith, H.E., Nematode sperm motility. WormBook, 2014: p. 1–15.

4. Roberts, T.M. and S. Ward, Centripetal flow of pseudopodial surface components could propel the amoeboid movement of Caenorhabditis elegans spermatozoa. J Cell Biol, 1982. 92(1): p. 132–8.

5. Nelson, G.A., T.M. Roberts, and S. Ward, Caenorhabditis elegans spermatozoan locomotion: amoeboid movement with almost no actin. J Cell Biol, 1982. 92(1): p. 121–31.

6. Miller, M.A., et al., A sperm cytoskeletal protein that signals oocyte meiotic maturation and ovulation. Science, 2001. 291(5511): p. 2144–7.

7. Govindan, J.A., et al., Somatic cAMP signaling regulates MSP-dependent oocyte growth and meiotic maturation in C. elegans. Development, 2009. 136(13): p. 2211–21.

8. Burke, D.J. and S. Ward, Identification of a large multigene family encoding the major sperm protein of Caenorhabditis elegans. J Mol Biol, 1983. 171(1): p. 1–29.

9. Scott, A.L., et al., Major sperm protein genes from Onchocerca volvulus. Mol Biochem Parasitol, 1989. 36(2): p. 119–26.

10. Scott, A.L., Nematode sperm. Parasitol Today, 1996. 12(11): p. 425–30.

11. Stanfield, G.M. and A.M. Villeneuve, Regulation of sperm activation by SWM-1 is required for reproductive success of C. elegans males. Curr Biol, 2006. 16(3): p. 252–63.

12. Roberts, T.M., F.M. Pavalko, and S. Ward, Membrane and cytoplasmic proteins are transported in the same organelle complex during nematode spermatogenesis. J Cell Biol, 1986. 102(5): p. 1787–96.

13. Yushin, V.V., M. Claeys, and W. Bert, Ultrastructural immunogold localization of major sperm protein (MSP) in spermatogenic cells of the nematode Acrobeles complexus (Nematoda, Rhabditida). Micron, 2016. 89: p. 43–55.

14. Zhu, G.D. and S.W. L’Hernault, The Caenorhabditis elegans spe-39 gene is required for intracellular membrane reorganization during spermatogenesis. Genetics, 2003. 165(1): p. 145–57.

15. Zhu, G.D., et al., SPE-39 family proteins interact with the HOPS complex and function in lysosomal delivery. Mol Biol Cell, 2009. 20(4): p. 1223–40.

16. Arduengo, P.M., et al., The presenilin protein family member SPE-4 localizes to an ER/Golgi derived organelle and is required for proper cytoplasmic partitioning during Caenorhabditis elegans spermatogenesis. J Cell Sci, 1998. 111 ( **Pt 24**): p. 3645–54.

17. Price, K.L., et al., The intrinsically disordered protein SPE-18 promotes localized assembly of MSP in Caenorhabditis elegans spermatocytes. Development, 2021. 148(5).

18. Liu, Z., et al., The micronutrient element zinc modulates sperm activation through the SPE-8 pathway in Caenorhabditis elegans. Development, 2013. 140(10): p. 2103–7.

19. Zhao, Y., et al., The zinc transporter ZIPT-7.1 regulates sperm activation in nematodes. PLoS Biol, 2018. 16(6): p. e2005069.

20. Roberts, T.M. and M. Stewart, Acting like actin. The dynamics of the nematode major sperm protein (msp) cytoskeleton indicate a push-pull mechanism for amoeboid cell motility. J Cell Biol, 2000. 149(1): p. 7–12.

21. Dickinson, R.B. and D.L. Purich, Nematode sperm motility: nonpolar filament polymerization mediated by end-tracking motors. Biophys J, 2007. 92(2): p. 622–31.

22. Mei, X., et al., SPE-51, a sperm-secreted protein with an immunoglobulin-like domain, is required for fertilization in C. elegans. Curr Biol, 2023. 33(14): p. 3048–3055 e6.

23. Varkey, J.P., et al., The Caenorhabditis elegans spe-6 gene is required for major sperm protein assembly and shows second site non-complementation with an unlinked deficiency. Genetics, 1993. 133(1): p. 79–86.

24. Songyang, Z., et al., Recognition of unique carboxyl-terminal motifs by distinct PDZ domains. Science, 1997. 275(5296): p. 73–7.

25. Harris, B.Z. and W.A. Lim, Mechanism and role of PDZ domains in signaling complex assembly. J Cell Sci, 2001. 114(Pt 18): p. 3219–31.

26. Ackermann, F., et al., The Multi-PDZ domain protein MUPP1 as a lipid raft-associated scaffolding protein controlling the acrosome reaction in mammalian spermatozoa. J Cell Physiol, 2008. 214(3): p. 757–68.

27. Irino, Y., et al., Phospholipase Cdelta4 associates with glutamate receptor interacting protein 1 in testis. J Biochem, 2005. 138(4): p. 451–6.

28. Jeong, M.G., et al., Transcriptional coactivator with PDZ-binding motif is required to sustain testicular function on aging. Aging Cell, 2017. 16(5): p. 1035–1042.

29. Zhang, Y.L. and Z.F. Han, Rational design of an orthogonal noncovalent interaction system at the MUPP1 PDZ11 complex interface with CaMKIIalpha-derived peptides in human fertilization. Mol Biosyst, 2017. 13(10): p. 2145–2151.

30. Liang, A.J., et al., The expression of the new epididymal luminal protein of PDZ domain containing 1 is decreased in asthenozoospermia. Asian J Androl, 2018. 20(2): p. 154–159.

31. Chu, D.S., et al., Sperm chromatin proteomics identifies evolutionarily conserved fertility factors. Nature, 2006. 443(7107): p. 101–5.

32. Brenner, S., The genetics of Caenorhabditis elegans. Genetics, 1974. 77(1): p. 71–94.

33. Dokshin, G.A., et al., Robust Genome Editing with Short Single-Stranded and Long, Partially Single-Stranded DNA Donors in Caenorhabditis elegans. Genetics, 2018. 210(3): p. 781–787.

34. Dickinson, D.J., et al., Streamlined Genome Engineering with a Self-Excising Drug Selection Cassette. Genetics, 2015. 200(4): p. 1035–49.

35. Wu, J.C., et al., Sperm development and motility are regulated by PP1 phosphatases in Caenorhabditis elegans. Genetics, 2012. 190(1): p. 143–57.

36. Chen, S.Y., et al., C. elegans spermatocyte divisions show a weak spindle checkpoint response. J Cell Sci, 2024. 137(6).

37. Shakes, D.C., et al., Spermatogenesis-specific features of the meiotic program in Caenorhabditis elegans. PLoS Genet, 2009. 5(8): p. e1000611.

38. Hsu, J.Y., et al., Mitotic phosphorylation of histone H3 is governed by Ipl1/aurora kinase and Glc7/PP1 phosphatase in budding yeast and nematodes. Cell, 2000. 102(3): p. 279–91.

39. Nishimura, H. and S.W. L’Hernault, Spermatogenesis-defective (spe) mutants of the nematode Caenorhabditis elegans provide clues to solve the puzzle of male germline functions during reproduction. Dev Dyn, 2010. 239(5): p. 1502–14.

40. Merta, H., et al., Cell cycle regulation of ER membrane biogenesis protects against chromosome missegregation. Dev Cell, 2021. 56(24): p. 3364–3379 e10.

41. Ferrandiz, N., et al., Endomembranes promote chromosome missegregation by ensheathing misaligned chromosomes. J Cell Biol, 2022. 221(6).

42. Grant, R.P., et al., Structure of MFP2 and its function in enhancing MSP polymerization in Ascaris sperm amoeboid motility. J Mol Biol, 2005. 347(3): p. 583–95.

43. Morrison, K.N., et al., MFP1/MSD-1 and MFP2/NSPH-2 co-localize with MSP during C. elegans spermatogenesis. MicroPubl Biol, 2021. 2021.

44. Lee, H.J. and J.J. Zheng, PDZ domains and their binding partners: structure, specificity, and modification. Cell Commun Signal, 2010. 8: p. 8.

45. Kuriu, T., et al., Differential control of postsynaptic density scaffolds via actin-dependent and -independent mechanisms. J Neurosci, 2006. 26(29): p. 7693–706.

46. Su, W.H., et al., Polarity protein complex Scribble/Lgl/Dlg and epithelial cell barriers. Adv Exp Med Biol, 2012. 763: p. 149–70.

47. Bonello, T.T., W. Choi, and M. Peifer, Scribble and Discs-large direct initial assembly and positioning of adherens junctions during the establishment of apical-basal polarity. Development, 2019. 146(22).

48. Gao, Y., et al., Cell polarity proteins and spermatogenesis. Semin Cell Dev Biol, 2016. 59: p. 62–70.

49. Wong, E.W., et al., Par3/Par6 polarity complex coordinates apical ectoplasmic specialization and blood-testis barrier restructuring during spermatogenesis. Proc Natl Acad Sci U S A, 2008. 105(28): p. 9657–62.

50. Heydecke, D., et al., The multi PDZ domain protein MUPP1 as a putative scaffolding protein for organizing signaling complexes in the acrosome of mammalian spermatozoa. J Androl, 2006. 27(3): p. 390–404.

51. Mruk, D.D. and C.Y. Cheng, The Mammalian Blood-Testis Barrier: Its Biology and Regulation. Endocr Rev, 2015. 36(5): p. 564–91.

